# Chromosome biorientation requires Aurora B’s spatial separation from its outer kinetochore substrates but not its turnover at kinetochores

**DOI:** 10.1101/2023.02.26.530110

**Authors:** Shuyu Li, Luis J. Garcia-Rodriguez, Tomoyuki U. Tanaka

**Author notes:** Contributed equally to the work (Shuyu Li, Luis J. Garcia-Rodriguez).

## Abstract

For correct chromosome segregation in mitosis, sister kinetochores must interact with microtubules from opposite spindle poles (biorientation). For this, aberrant kinetochore– microtubule interaction must be resolved (error correction) by Aurora B kinase. Once biorientation is formed, tension is applied across sister kinetochores, stabilizing kinetochore– microtubule interactions. The mechanism for this tension-dependent process has been debated. Here we study how localizations of Aurora B at different kinetochore sites affect the establishment and maintenance of biorientation in budding yeast. In the absence of the physiological Aurora B–INCENP recruitment mechanisms, engineered recruitment of Aurora B–INCENP to the inner kinetochore (Mif2) prior to biorientation supports the subsequent establishment of biorientation. By contrast, an engineered Aurora B–INCENP recruitment to the outer kinetochore (Ndc80) fails to support biorientation establishment. Furthermore, when the physiological Aurora B–INCENP recruitment mechanisms are present, an engineered Aurora B–INCENP recruitment to Mif2 during metaphase (after biorientation establishment) does not affect biorientation maintenance. By contrast, an engineered Aurora B–INCENP recruitment to Ndc80 during metaphase leads to disruption of biorientation, which is dependent on the kinase activity of Aurora B. Taken together, our results suggest that spatial separation of Aurora B from its outer kinetochore substrates is required to stabilize kinetochore–microtubule interaction when biorientation is formed and tension is applied on this interaction. Meanwhile, Aurora B shows dynamic turnover (or exchange) on the centromere and kinetochore during early mitosis. It has been thought that this turnover is crucial for error correction and biorientation, as it may help Aurora B reach its substrates in distance and/or may facilitate the Aurora B activation on the mitotic spindle. However, using the engineered Aurora B–INCENP recruitment to the inner kinetochore, we demonstrate that, even without such a turnover, Aurora B–INCENP can efficiently support biorientation. Altogether, our study provides important insights into how Aurora B promotes error correction and biorientation in a tension-dependent manner.

## Introduction

To ensure correct chromosome segregation in mitosis, sister kinetochores must interact with microtubules (MTs) extending from opposite spindle poles (chromosome biorientation). To establish biorientation, any aberrant kinetochore–MT interactions (e.g. syntelic attachment where both sister kinetochores interact with MTs from the same spindle pole) must be weakened and disrupted in the process called error correction [1-4]. This process relies on the phosphorylation of outer kinetochore components by Aurora B kinase (Ipl1 in budding yeast) [5-7]. In budding yeast, the Dam1 complex is the most important Aurora B substrate for the disruption of aberrant kinetochore–MT interaction, while phosphorylations of the N-terminus of Ndc80 (a component of the Ndc80 complex; Ndc80C) modestly contribute to the process [8, 9] (Figure 1A, top). The disruption of aberrant kinetochore–MT interaction is followed by new kinetochore–MT interaction [10, 11]. If this new interaction leads to the establishment of chromosome biorientation, tension is applied across sister kinetochores, which stabilizes kinetochore–MT interactions. It has long been debated how Aurora B stops disrupting kinetochore–MT interaction when tension is applied.

**Figure 1.**
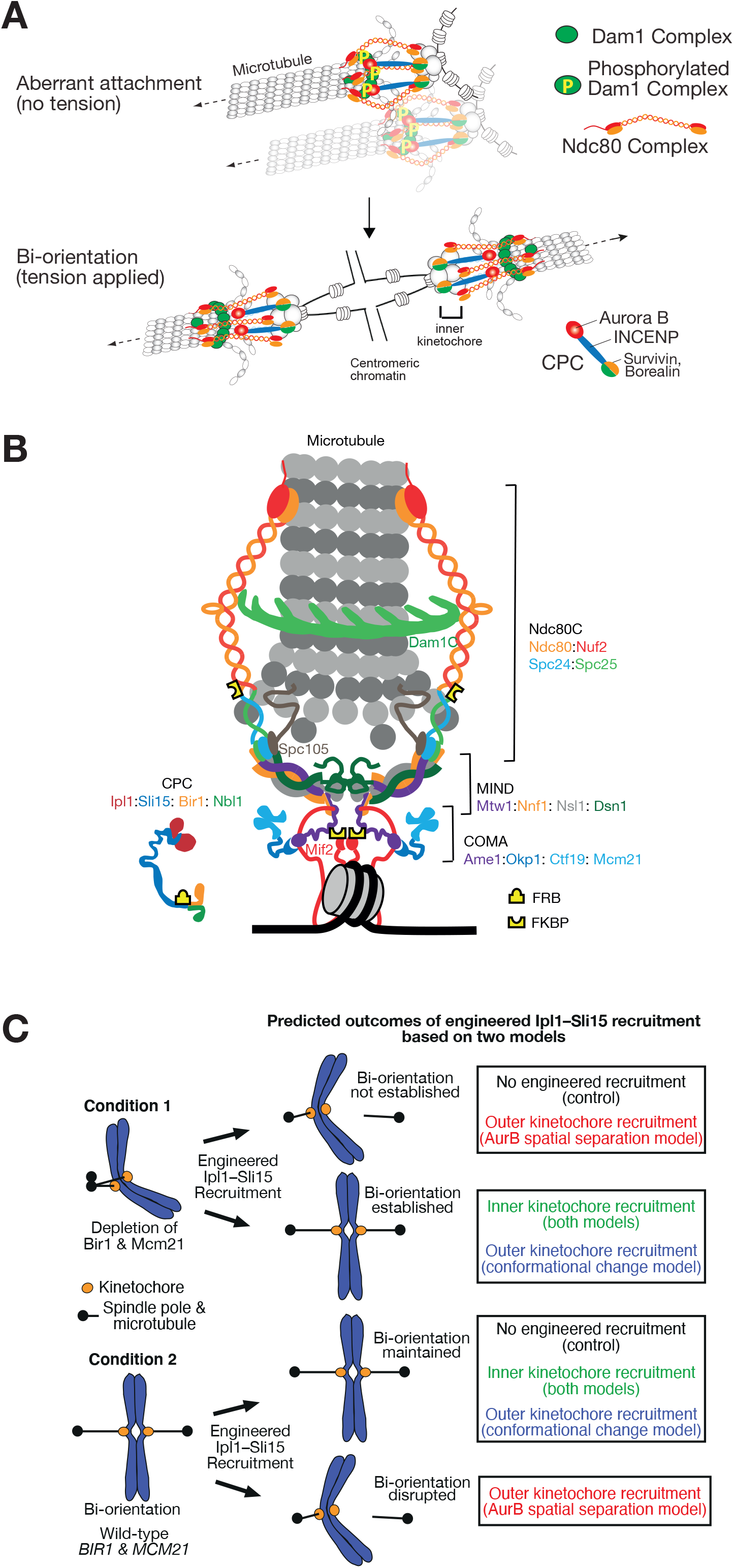
Introductory diagrams and the design of experiments. **A**. Diagram illustrating aberrant (syntelic) kinetochore–MT interactions (top) and chromosome biorientation (bottom) in budding yeast. To resolve aberrant kinetochore-MT interaction, the Dam1-complex components must be phosphorylated by Aurora B kinase (Ipl1 in budding yeast), which weakens and disrupts kinetochore–MT interaction (top). When biorientation is established and tension is applied across sister kinetochores, kinetochore–MT interactions are stabilized (bottom). According to the Aurora B spatial separation model, the Ndc80 complex is stretched when tension is applied, leading to spatial separation of Aurora B from the Dam1 complex. See references and more details in text. **B**. Diagram illustrating the kinetochore structure in budding yeast. Protein complexes making and regulating the kinetochore (Dam1C, Ndc80C, MIND, COMA and CPC) and their components are shown in colors. In the current study, FRB was fused to the N-terminus of Sli15, and FKBP12 to the C-terminus of Mif2 or Ndc80 (highlighted in yellow). This diagram is based on those in [23, 24] but contains some modifications. **C**. Diagram showing the design of experiments in this study. We test whether the engineered Ipl1–Sli15 recruitment to the inner or outer kinetochore i) supports biorientation establishment in the absence of physiological Ipl1–Sli15 recruitment mechanisms (condition 1) and ii) supports (or disrupts) biorientation maintenance in the presence of physiological Ipl1–Sli15 recruitment mechanisms (condition 2). Predicted outcomes are shown in colors within rectangles in the right, based on the Aurora B (AurB) spatial separation model and the kinetochore conformational change model (see text).

To understand this mechanism, we need to consider where Aurora B localizes at the centromere and kinetochore. Aurora B forms the Chromosomal Passenger Complex (CPC) together with INCENP, Survivin and Borealin (Sli15, Bir1, and Nbl1, respectively, in budding yeast) [12]. Several studies reported that the CPC is recruited to the centromere, which is mediated by the interaction of Survivin with Shugoshin and phosphorylated histones [13-17]. However, more recently, a Survivin (Bir1)-independent CPC recruitment mechanism was found in budding yeast. In this mechanism, INCENP (Sli15) directly interacts with the Mcm21– Ctf19 subcomplex at the inner kinetochore for recruitment of the CPC [18, 19]. The two mechanisms for CPC recruitment, i.e. Survivin (Bir1)-dependent centromere recruitment and Survivin (Bir1)-independent inner kinetochore recruitment, work redundantly to promote chromosome biorientation in budding yeast. If both mechanisms become defective, most sister kinetochores fail to establish biorientation.

To explain how Aurora B stops disrupting kinetochore–MT interaction when tension is applied, several models have been proposed [4, 20]. In the current study, we focus on the following two models: First, when tension is applied, the long coiled-coil region of the Ndc80C is stretched and exceeds INCENP in length, which would spatially separate Aurora B from its outer kinetochore substrates whose phosphorylation is crucial for error correction [3, 5, 21, 22] (Figure 1A, bottom). The spatial separation would happen in budding yeast because, when bi-orientation is established, the centromere and the inner kinetochore (where the CPC is recruited) are adjacent to each other (<10–20 nm) but apart from the outer kinetochore substrates of Aurora B (> 60 nm) [23, 24] (Figure 1B). The spatial separation would lead to dephosphorylation of the outer kinetochore substrates [25, 26], stabilizing the kinetochore attachment to the MT end (Aurora B spatial separation model). On the other hand, when tension is low, a kink in the middle of Ndc80C allows the Ndc80C to bend flexibly [27-29], which would enable Aurora B to access its outer kinetochore substrates (Figure 1A, top). Second, when tension is applied to kinetochore–MT interaction, the kinetochore might undergo a conformational change in such a way that the kinetochore attachment to the MT end is stabilized (kinetochore conformational change model) [30]. Such stabilization would happen if the kinetochore (or CPC) conformational change either a) leads to dephosphorylation of the outer kinetochore substrates of Aurora B or b) overcomes the effect of Aurora B-dependent phosphorylation on weakening the kinetochore attachment to the MT end.

One way to test the Aurora B spatial separation model would be to assess the establishment and maintenance of biorientation after ectopically targeting Aurora B (with INCENP that is required for Aurora B kinase activity) to different sites within the kinetochore. Such targeting should be carried out to compare the outcomes, in the presence and absence of physiological mechanisms for Aurora B recruitment to the centromere/kinetochore. In their absence, we will understand the location of Aurora B that suffices its role for error correction, while in their presence we will address the dominant effects of Aurora B in destabilizing kinetochore–MT interaction. Meanwhile, the evidence for the kinetochore conformational change model was recently obtained *in vitro* using purified yeast kinetochores [30]. However, this model should be tested further *in vivo* in cells.

Here, to test these models, we engineered recruitment of the Aurora B–INCENP (Ipl1–Sli15 in yeast) to the inner kinetochore and outer kinetochore in budding yeast cells, with and without the physiological Aurora B recruitment mechanisms. We also tested two different orientations of the INCENP for the engineered recruitment. Our results suggest that a) the Aurora B spatial separation from its outer kinetochore substrates is required for chromosome biorientation and b) the kinetochore conformational change alone is not sufficient to promote biorientation. Meanwhile, the CPC shows dynamic turnover on the centromere and kinetochore during early mitosis [31-33], and this turnover is possibly required for error correction as it may help Aurora B reach its substrates in distance and/or may facilitate the Aurora B activation on the mitotic spindle [20, 34]. Using the engineered CPC recruitment, we also test whether the dynamic turnover of the CPC at the kinetochore is required for error correction.

## Results

### Designing engineered Ipl1–Sli15 recruitment to the kinetochore to test the Aurora B spatial separation model vs the kinetochore conformational change model

We intended to test whether the Aurora B spatial separation or the kinetochore conformational change is the major mechanism to stabilize chromosome biorientation in budding yeast cells. For this, we engineered recruitment of Ipl1–Sli15 to the inner kinetochore and the outer kinetochore. We used FRB and FKBP protein domains, which form a heterodimer in the presence of Rapamycin [35]. FRB was fused to the N-terminus of Sli15 (FRB-Sli15) while FKBP12 was fused to the C-terminus of Mif2 (yeast orthologue of CENP-C, localizing at the inner kinetochore) or Ndc80 (localizing at the outer kinetochore) [23] (Figure 1B). We carried out this engineered recruitment in two conditions: First, we depleted Bir1 and Mcm21 using an auxin-dependent degron (Bir1-aid, Mcm21-aid) when cells were released from the arrest at G1 phase, which abolished physiological mechanisms of Ipl1–Sli15 recruitment to the inner kinetochore/centromere [18, 19]. We then added Rapamycin to induce the engineered recruitment, prior to the establishment of biorientation and observed whether biorientation is subsequently established or not (Figure 1C, condition 1). Note that Mcm21 depletion does not affect Mif2 or the outer kinetochore assembly [36] and therefore should not affect the engineered Ipl1–Sli15 recruitment. Second, we allowed cells to establish biorientation with intact physiological Ipl1–Sli15 recruitment mechanisms (i.e. with wild-type Bir1 and Mcm21). We then added Rapamycin to induce the engineered recruitment during metaphase arrest (after the establishment of biorientation) and addressed whether biorientation was maintained or disrupted (Figure 1C, condition 2).

The predicted outcomes of these experiments are as follows: The engineered Ipl1-Sli15 recruitment to the outer kinetochore would not allow spatial separation of Ipl1 from its outer kinetochore substrates when tension is applied across sister kinetochores. Therefore, if the Ipl1 spatial separation is required to stabilize biorientation when tension is applied, biorientation would not be stably established in condition 1, and biorientation would be disrupted in condition 2 (Figure 1C, highlighted in red). On the other hand, the engineered Ipl1 recruitment to the inner kinetochore would allow spatial separation of Ipl1 from its outer kinetochore substrates when tension is applied. Therefore, even if the Ipl1 spatial separation is required to stabilize biorientation, biorientation would be stably established in condition 1, and biorientation would be maintained in condition 2 (Figure 1C, highlighted in green). Meanwhile, if the kinetochore conformational change is the mechanism to stabilize biorientation when tension is applied, Ipl1 at either the inner or outer kinetochore would promote error correction (as far as it destabilizes kinetochore–MT interaction in the absence of tension) and would not disrupt already-established biorientation. Therefore, biorientation would be stably established in condition 1 and would be maintained in condition 2 with engineered Ipl1-Sli15 recruitment either to the outer kinetochore (Figure 1C, highlighted in blue) or inner kinetochore (highlighted in green).

To interpret the outcomes from these experiments, we need evidence that Ipl1 in the engineered recruitment is indeed functional to change kinetochore–MT interaction under a suitable condition (e.g. with low tension). This would not be the case, for example, if Ipl1 cannot be activated, is sterically interfered with, or is oriented away from its outer kinetochore substrates in the engineered recruitment. In these situations, the engineered recruitment would make no difference from the control (no engineered Ipl1-Sli15 recruitment; Figure 1C, black in rectangles) in condition 1 or 2. We can test this by conducting experiments in both conditions 1 and 2, with the outer or inner kinetochore recruitment of CPC. In fact, in either condition 1 or 2, the predicted outcome based on either model (Aurora B spatial separation model or kinetochore conformational model) would be different from the control (Figure 1C, compare each red, blue or green with the control [black], in rectangles). For example, with the Aurora B spatial separation model, the outer kinetochore recruitment would show a different outcome from the control in condition 2, (Figure 1C, compare red with the control [black] in rectangles). Therefore, if outcomes fit the predictions from either model (Figure 1C, red, blue or green in rectangles), we also confirm that Ipl1 is functional in the engineered recruitment.

### The engineered recruitment of Ipl1–Sli15 to the inner and outer kinetochore in the absence of Bir1 restores its physiological level at the kinetochore

Before conducting experiments in conditions 1 and 2 (Figure 1C), we addressed if the engineered recruitment of Ipl1–Sli15 worked as expected at the inner and outer kinetochore. To distinguish its localization at the kinetochore from that at spindle MTs, we isolated a chosen centromere, *CEN3*, from the mitotic spindle by inactivating *CEN3* and thereby inhibiting its kinetochore–MT interaction [37, 38] (Figure 2A, left). Subsequently, we reactivated *CEN3* to allow kinetochore assembly on it. We then evaluated the amount of Ipl1 (visualized with GFP fusion) at the kinetochore (on *CEN3*) before it interacted again with spindle MTs (Figure 2A, right).

**Figure 2.**
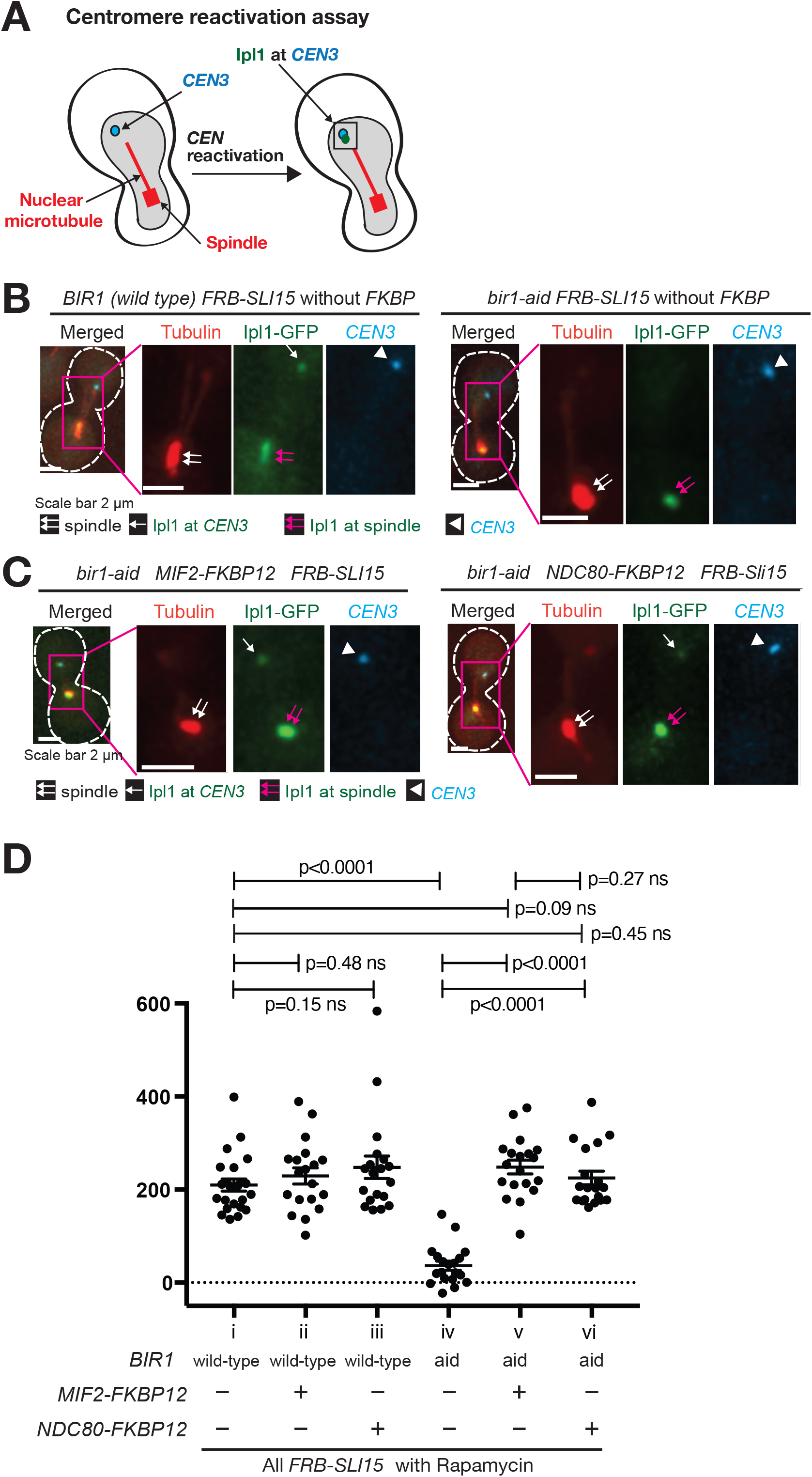
The engineered Ipl1–Sli15 recruitment to the inner and outer kinetochore in the absence of Bir1 restores its physiological level at the kinetochore. **A**. The diagram shows the method of quantifying Ipl1 signals at an isolated *CEN3. CEN3* under *GAL1-10* promoter was inactivated by transcription from the promoter, which prevented its interaction with MTs and placed it away from the spindle (left) [37, 38]. After reactivation of *CEN3* (by shutting off the promoter) during metaphase arrest, Ipl1 at *CEN3* was quantified (right). **B.C. D**. FKBP12-fused Mif2 and FKBP12-fused Ndc80 recruit Ipl1-GFP (bound by FRB-fused Sli15) to an isolated *CEN3*. Six strains contained the combination of i) *BIR1+* (wile-type) or *bir1-aid* and ii) no *FKBP12, MIF2*-*FKBP12* or *NDC80-FKBP12*, as shown below in graph **D** (T13985, T13982, T13979, T13983, T13984 and T13981 from left to right). All six strains also contained *FRB-SLI15, IPL1-GFP, TIR, GAL1-10* promoter*-CEN3-tetOs, TetR-3xCFP, mCherry-TUB1* and *MET3* promoter*-CDC20*. Cells were released from G1 arrest and subsequently arrested in metaphase by Cdc20 depletion while *CEN3* was inactivated (**A**, left). Immediately after *CEN3* was reactivated during metaphase arrest (**A**, right), fluorescence images were acquired for 10 min with a 1-min interval. 500 µM NAA was added (to deplete Bir1-aid if present) 30min before the release from G1 arrest, and 1 µM rapamycin was added (to induce FRB-FKBP12 dimerization if they are present) 30 mins before *CEN3* reactivation. Representative cells (T13985 [**B**, left], T13983 [**B**, right], T13984 [**C**, left], T13981 [**C**, right]) are shown. Cell shapes are shown in white broken lines. Ipl1-GFP signals associated at *CEN3* for the last four time points prior to *CEN3* reaching the spindle (or for the last four time points of the observation if *CEN3* did not reach the spindle during the observation) were quantified, added and shown in graph **D**, with mean and error bars (SEM). *p* values were obtained by t-test.

In the presence of FRB-Sli15, we compared the level of Ipl1-GFP at *CEN3* with and without FKBP12 fusion to Mif2 (or Ndc80) and with and without depletion of Bir1 (Figure 2B–D). Rapamycin was added in all conditions to induce dimerization between FRB and FKBP12 if they were present. Auxin (NAA) was also added in all conditions to induce Bir1 depletion if *bir1-aid* was present. When FKBP12 was not fused to Mif2 (or Ndc80), Bir1 depletion considerably reduced the level of Ipl1 at *CEN3* (Figure 2B; 2D, compare i and iv) which is consistent with our previous result [18]. The fusion of FKBP12 to Mif2 restored Ipl1 localization at *CEN3* to the normal level (Figure 2C, left; 2D, compare v with i and iv). Thus, Ipl1–Sli15 was indeed recruited to the kinetochore by the inner kinetochore component Mif2 in the engineered system. Similarly, the fusion of FKBP12 to Ndc80 restored Ipl1 localization at *CEN3* to the normal level (Figure 2C, right; 2D, compare vi with i and iv). Thus, Ipl1–Sli15 was also recruited to the kinetochore by the outer kinetochore component Ndc80 in the engineered system.

### The engineered recruitment of Ipl1–Sli15 to the inner kinetochore, but not to the outer kinetochore, facilitates biorientation establishment in the absence of its physiological recruitment

We then conducted experiments in condition 1, as planned in Figure 1C (top). We depleted Bir1-aid and Mcm21-aid by addition of auxin (NAA) when cells were released from the arrest at G1 phase in order to abolish physiological mechanisms of Ipl1–Sli15 recruitment to the inner kinetochore/centromere [18, 19]. At the same time, we also induced interaction between FRB-Sli15 and either Mif2-FKBP12 or Ndc80-FKBP12 by addition of Rapamycin, prior to the establishment of biorientation (Figure 3A, B).

**Figure 3.**
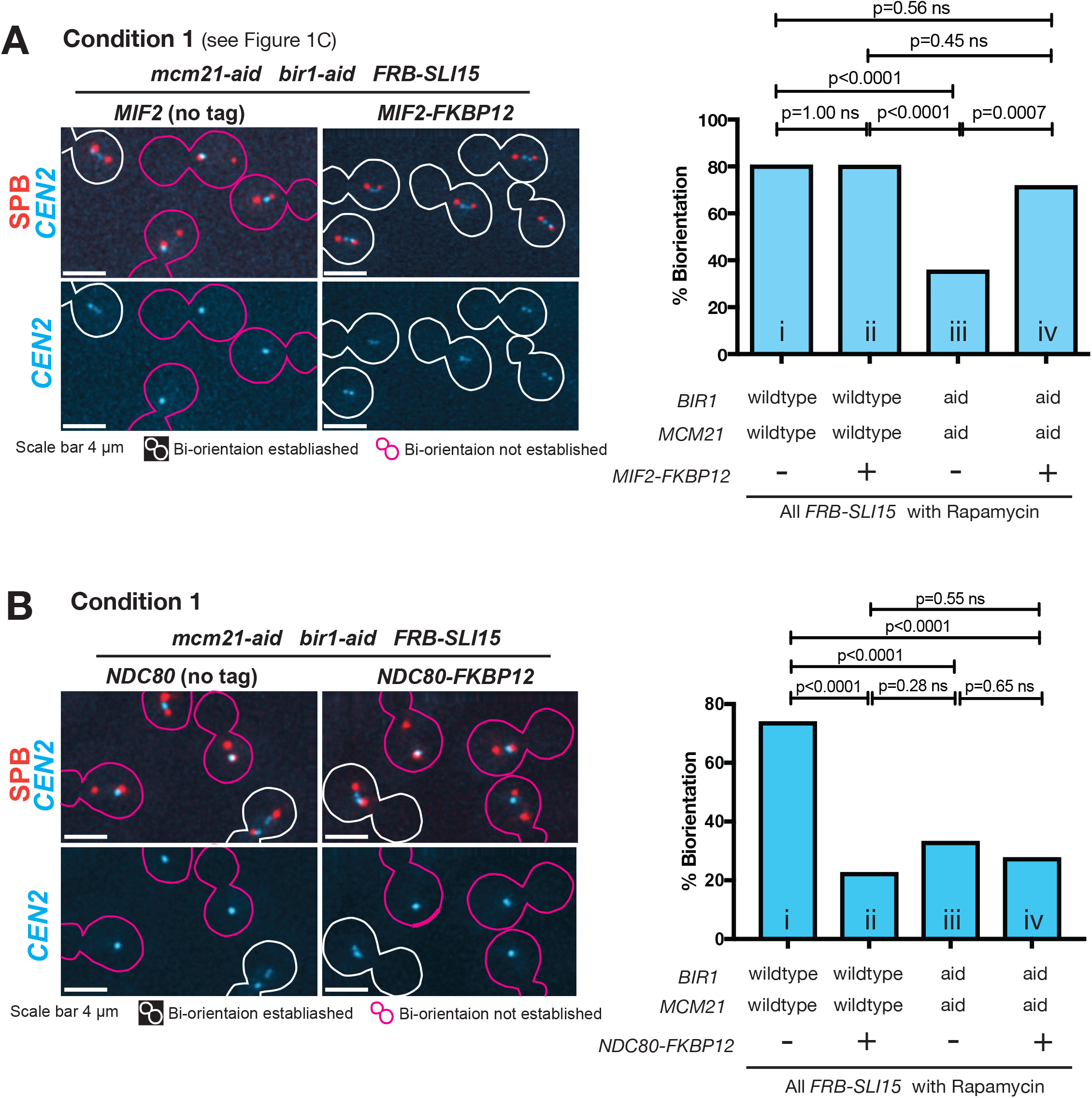
The engineered Ipl1–Sli15 recruitment to the inner kinetochore, but not the outer kinetochore, restores bi-orientation when its physiological recruitment mechanisms are defective. **A**. The engineered Ipl1–Sli15 recruitment to Mif2 facilitated bi-orientation establishment after Bir1 and Mcm22 were depleted. Four strains contained the combination of i) *BIR1+ MCM21+* (wile-type) or *bir1-aid mcm21-aid* and ii) *MIF2* with or without fusion to *FKBP12*, as shown below in the graph (T13942, T13946, T13943 and T13945 from left to right). All four strains also carried *FRB-SLI15, TIR, CEN2-tetOs, TetR-3xCFP, SPC42-4xmCherry, and MET3 promoter-CDC20*. Cells of these strains were released from G1 arrest and subsequently arrested in metaphase by Cdc20 depletion. From 2 h following the release from G1 arrest, fluorescence images were acquired for 5 min with 30 s intervals. 500 µM NAA was added (to deplete Bir1-aid and Mcm21-aid if present) 30min before the release from G1 arrest, and 1 µM rapamycin was added (to induce FRB-FKBP12 dimerization if they are present) 30 mins before the start of image acquisition. Representative cells (T13943 [left] and T13945 [right]) are shown on the left. Cell shapes are shown in white or magenta lines when biorientation was established or not established, respectively. The percentage of cells with biorientation is shown in the right graph (n=31, 72, 78 and 32 from the left to right). *p* values were obtained by Fisher’s exact test. **B**. The engineered Ipl1-Sli15 recruitment to Ndc80 did not restore biorientation after Bir1 and Mcm22 were depleted. Four strains contained the combination of i) *BIR1+ MCM21+* (wile-type) or *bir1-aid mcm21-aid* and ii) *NDC80* with or without fusion to *FKBP12*, as shown below in the graph (T13942, T13956, T13943 and T13964 from left to right). All four strains also carried the common constructs same as in **A**. Cells were treated as in **A**, and data were collected, analyzed and shown as in **A**. Representative cells (T13943 [left] and T13964 [right]) are shown on the left. The percentage of cells with biorientation is shown in the right graph (n=50, 63, 44 and 54 from the left to right).

In the control with wild-type Bir1 and Mcm21 (and without FKBP12 fusion), biorientation was established in most cells, i.e. sister *CEN2* dots separated on the metaphase spindle (Figure 3A, B; i in the graphs). The depletion of Bir1-aid and Mcm21-aid (without FKBP12 fusion) led to failure in biorientation establishment in the majority cells (Figure 3A, B; left images, iii in the graphs), as reported previously [18]. The engineered recruitment of Ipl1–Sli15 with Mif2-FKBP12 restored biorientation establishment to the normal level, despite the depletion of Bir1-aid and Mcm21-aid (Figure 3A; right images, iv in the graph). By contrast, the engineered recruitment of Ipl1–Sli15 to Ndc80-FKBP12 failed to restore biorientation establishment with depletion of Bir1-aid and Mcm21-aid (Figure 3B; right images, iv in the graph). With wild-type Bir1 and Mcm21, the engineered recruitment of Ipl1–Sli15 to Ndc80-FKBP12 also led to the failure of biorientation establishment in the majority of cells (Figure 3B, ii in the graph). However, this was not the case with the engineered recruitment of Ipl1–Sli15 to Mif2-FKBP12 (Figure 3A, ii in the graph). We conclude that the engineered recruitment of Ipl1–Sli15 to the inner kinetochore, but not to the outer kinetochore, restores biorientation establishment in the absence of its physiological recruitment mechanisms. This is consistent with the Aurora B spatial separation model, as shown for condition 1 in Figure 1C (top; see red and green in predictions within rectangles).

### The engineered recruitment of Ipl1–Sli15 to the outer kinetochore, but not to the inner kinetochore, during metaphase leads to disruption of chromosome bi-orientation

Next, we conducted experiments in condition 2, as planned in Figure 1C (bottom). We used cells with wild-type Bir1 and Mcm21, i.e. with intact physiological mechanisms of Ipl1-Sli15 recruitment to the kinetochore. These cells were released from G1 phase and arrested in metaphase. During metaphase arrest (i.e. after bi-orientation was established), we added Rapamycin to induce interaction between FRB-Sli15 and either Mif2-FKBP12 or Ndc80-FKBP12.

Before the addition of Rapamycin in metaphase, the cells established biorientation, i.e. sister *CEN2* dots showed separation, in most cells (Figure 4A, B; iii in the graphs). Critically, the engineered recruitment of Ipl1–Sli15 to Ndc80-FKBP12 by addition of Rapamycin, but not to Mif2-FKBP12, disrupted bi-orientation in majority of cells (Figure 4A, B; right images, iv in the graphs). This disruption was indeed due to Ipl1–Sli15 recruitment to Ndc80-FKBP12 because such disruption was not observed without FKBP12 fusion after the addition of Rapamycin (Figure 4B, left images; i, ii in the graph). It is unlikely that this disruption was caused by the potential reduction of Ipl1–Sli15 at the centromere/inner kinetochore due to its engineered recruitment to Ndc80 (this would happen if the total amount of Ipl1–Sli15 is rather limited) because, once biorientation is established, the loss of Ipl1 (at the centromere/inner kinetochore) does not lead to the disruption of biorientation [5]. We conclude that the engineered recruitment of Ipl1–Sli15 to the outer kinetochore, but not to the inner kinetochore, during metaphase leads to disruption of chromosome bi-orientation. So far, this is consistent with the Aurora B spatial separation model, as shown for condition 2 in Figure 1C (bottom; see red and green in predictions within rectangles). But we test this model further in the next section.

**Figure 4.**
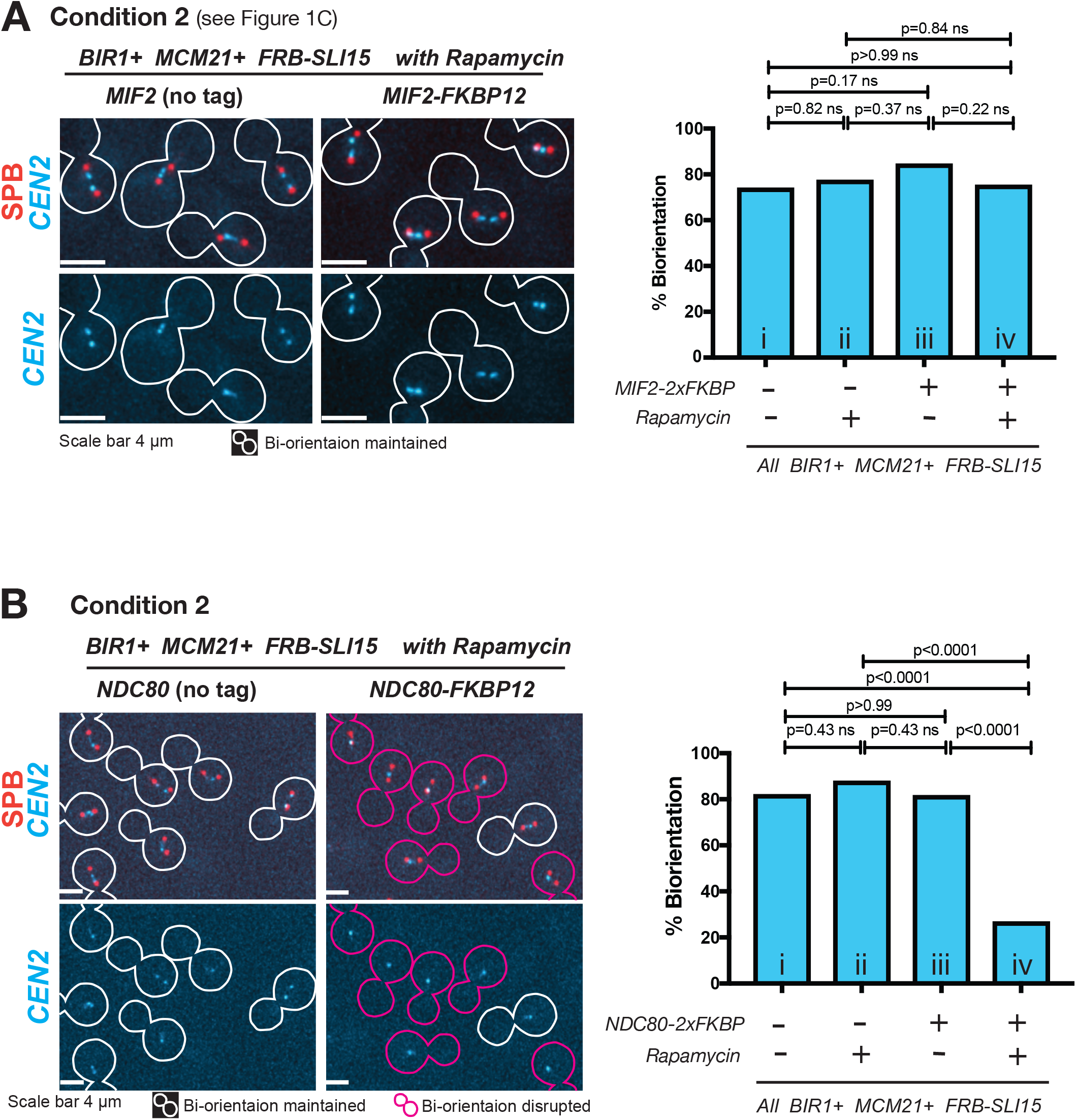
The engineered Ipl1–Sli15 recruitment to the outer kinetochore, but not to the inner kinetochore, during metaphase leads to disruption of biorientation, in the presence of physiological Ipl1–Sli15 recruitment mechanisms. **A**. The engineered Ipl1–Sli15 recruitment to the inner kinetochore during metaphase does not disrupt biorientation. Two *BIR1+ MCM21+* strains with and without *FKBP12* fusion to *MIF2* (T13946 and T13942, respectively) were analyzed. They also contained *FRB-SLI15, CEN2-tetOs, TetR-3xCFP, SPC42-4xmCherry, and MET3 promoter-CDC20*. Cells of the two strains were released from G1 arrest and subsequently arrested in metaphase by Cdc20 depletion. From 3 h following the release from G1 arrest, fluorescence images of an aliquot were acquired for 5 min with 30 s intervals (i and iii in the graph). At the same time, 1 µM rapamycin was added (to induce FRB-FKBP12 dimerization if they are present) to the original culture and, 1 h later, fluorescence images of a new aliquot were acquired in the same way (ii and iv in the graph). Representative cells are shown on the left. Cell shapes are shown in white lines when biorientation was maintained. The percentage of cells with biorientation is shown in the right graph (n=50, 62, 71 and 81 from left to right). *p* values were obtained by Fisher’s exact test. **B**. The engineered Ipl1–Sli15 recruitment to the outer kinetochore during metaphase leads to disruption of biorientation. Two *BIR1+ MCM21+* strains with and without *FKBP12* fusion to *NDC80* (T13956 and T13942, respectively) were analyzed. They also contained *FRB-SLI15, CEN2-tetOs, TetR-3xCFP, SPC42-4xmCherry, and MET3 promoter-CDC20*. Cells of the two strains were treated, imaged and analyzed as in **A**. Representative cells are shown on the left. Cell shapes are shown in white or magenta lines when biorientation was maintained or disrupted, respectively. The percentage of cells with biorientation is shown in the right graph (n=50, 66, 49 and 108 from left to right). *p* values were obtained by Fisher’s exact test.

### The disruption of bi-orientation by engineered Ipl1–Sli15 recruitment to the outer kinetochore requires Ipl1 kinase activity and is graded by different spatial arrangements of Ipl1–Sli15

We consider further how biorientation was disrupted by engineered Ipl1–Sli15 recruitment to the outer kinetochore in Figure 4B. The first possibility is that, because Ipl1 did not spatially separate from its outer kinetochore substrates, it continued phosphorylating them and disrupted kinetochore–MT interaction, as predicted by Aurora B spatial separation model. The second possibility is that this disruption was caused by spatial constraint, i.e. Ipl1–Sli15 recruited to the outer kinetochore may have physically interfered with kinetochore–MT interactions. The disruption of biorientation in the first possibility would be dependent on the Ipl1 kinase activity but the disruption in the second possibility would not. So, to test these possibilities, we used *ipl1-as5* mutant whose kinase activity is inhibited by the addition of an ATP analog 1NA-PP1 [39]. We conducted experiments in condition 2 with *ipl1-as5* (Figure 1C, bottom), i.e. after the establishment of biorientation, we added a) Rapamycin to induce Ipl1– Sli15 recruitment with Ndc80-FKBP12 and b) the ATP analog to inhibit *ipl1-as5* kinase activity. After such treatment, biorientation was still maintained with *ipl1-as5* (Figure 5A, right images, iv in the graph). By contrast, in the control with Ipl1 wild-type, the same treatment led to disruption of biorientation in the majority of cells (Figure 5A, left images, ii in the graph). This suggests that the disruption of biorientation by engineered Ipl1–Sli15 recruitment to the outer kinetochore in condition 2 (Figure 1C and 4B) is dependent on the Ipl1 kinase activity and therefore is not due to physical interference of kinetochore–MT interaction.

**Figure 5.**
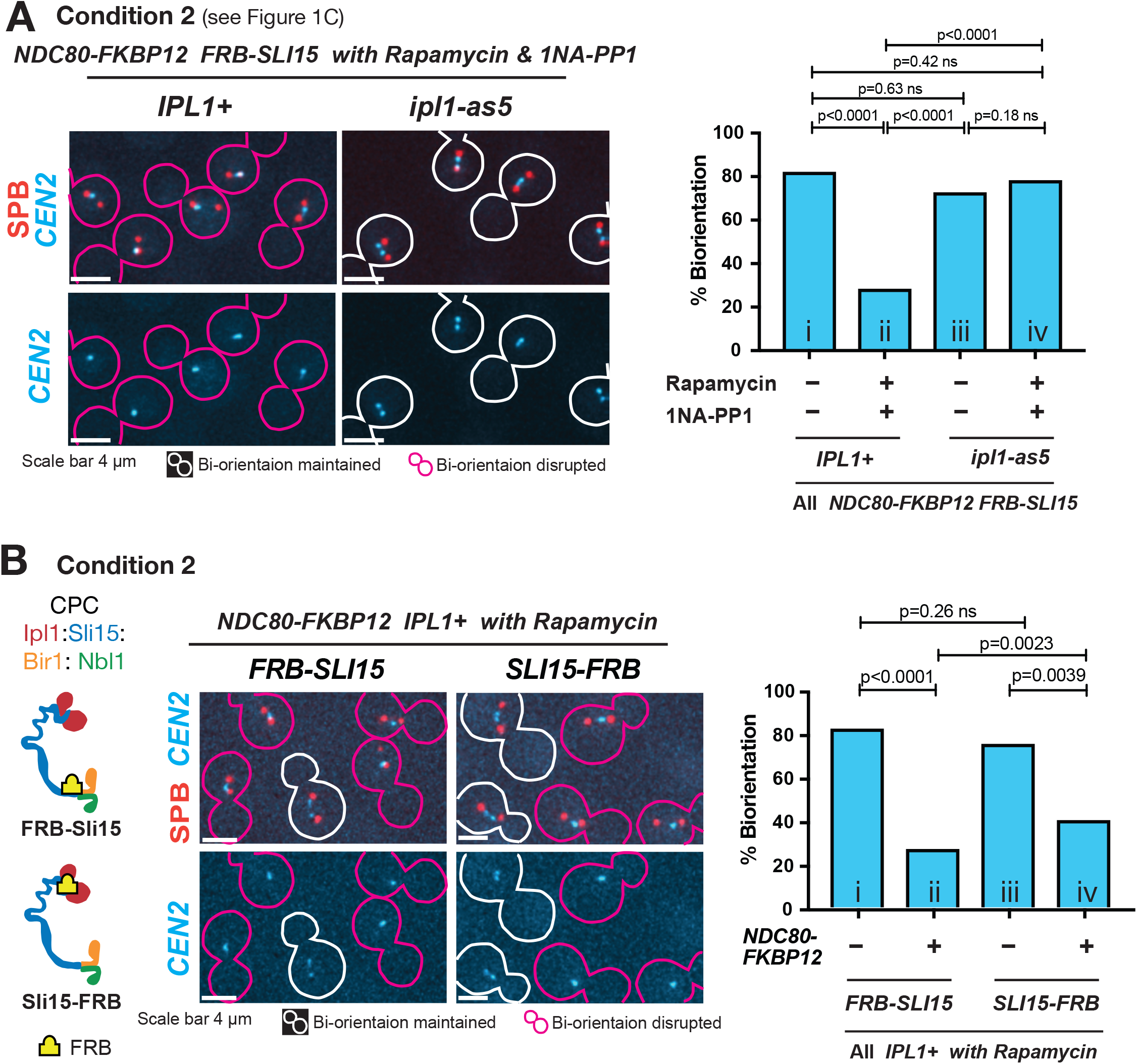
Ipl1 kinase activity was required for disruption of biorientation caused by engineered Ipl1-Sli15 recruitment to the outer kinetochore, and different spatial arrangements of Ipl1–Sli15 grade the disruption. **A**. Ipl1 kinase activity was required for disruption of biorientation caused by engineered Ipl1-Sli15 recruitment to the outer kinetochore during metaphase. Cells of *IPL1* wild-type and *ipl1-as5* mutant strains (T13976 and T13975, respectively), both carrying *FRB-SLI15, NDC80-2xFKBP, CEN2-tetOs, TetR-3xCFP, SPC42-4xmCherry, and MET3 promoter-CDC20* were released from G1 arrest and subsequently arrested in metaphase by Cdc20 depletion. At 3 h following the release from G1 arrest, 1 µM rapamycin (to induce FRB-FKBP12 dimerization) and 10 µM 1NA-PP1 (to inactivate Ipl1 kinase activity if *ipl1-as5* is present) were added to the media and incubated for 1 h. Fluorescence images were acquired for 5 min with 30-s intervals before and after adding rapamycin and 1NA-PP1. Representative cells are shown on the left. Cell shapes are shown in white or magenta lines when biorientation was maintained or disrupted, respectively. The percentage of cells with biorientation is shown in the right graph (n=56, 57, 46 and 54 from left to right). *p* values were obtained by Fisher’s exact test. **B**. FRB fusion to the C-terminus of Sli15 shows a more modest disruption of biorientation than does FRB fusion to the N-terminus of Sli15. Diagrams on the left show the position of FRB fusion to Sli15. Cells of four strains carrying *FRB-SLI15* (T13942), *FRB-SLI15 NDC80-2xFKBP* (T13956), *SLI15-FRB* (T13434) and *SLI15-FRB, NDC80-2FKBP* (T13795) were treated and analyzed as the Figure 4, except that NAA was not added. Representative cells are shown in the middle. The percentage of cells with biorientation is shown in the right graph (n=38, 108, 66 and 70). *p* values were obtained using Fisher’s exact test.

If the Aurora B spatial separation model is correct, we envisage that the disruption of biorientation may be graded by changing the spatial position of Ipl1 relative to its outer kinetochore substrates (Dam1C and the Ndc80 N-terminus). To test this, we fused FRB to the C-terminus of Sli15 (Sli15-FRB), instead of the N-terminus (FRB-Sli15), and analysed the outcome of Ipl1–Sli15 recruitment to the outer kinetochore (to Ndc80-FKBP12). Because Sli15 is an extended protein whose C-terminus binds and activates Ipl1 [40], we expect that Sli15-FRB would position Ipl1 in the vicinity of the Ndc80 C-terminus, i.e. somewhat further away from the outer kinetochore substrates of Ipl1 than would FRB-Sli15 do (Figure 5B, diagrams in left; also refer to Figure 1B). Similarly to FRB-Sli15, Sli15-FRB restored the normal level of Ipl1–Sli15 on an isolated kinetochore, when recruited to Ndc80-FKBP12 after Bir1 depletion, in an experiment similar to Figure 2C, D (Figure S1). We then conducted experiments in condition 2 (Figure 1C, bottom) with FRB-Sli15 and Sli15-FRB – we induced their recruitment to Ndc80-FKBP12 after biorientation was established. Crucially, compared with FRB-Sli15, Sli15-FRB showed a reduced level of biorientation disruption, (Figure 5B; compare left and right in images, and ii and iv in graph). Thus, the disruption of biorientation was graded by different spatial arrangements of Ipl1 in the engineered recruitment, as predicted from the Aurora B spatial separation model. The results in Figure 5A and 5B provide further evidence that the Aurora B spatial separation from its outer kinetochore substrates is required to stabilize kinetochore–MT interaction when biorientation is established.

### Evidence that rapid turnover of Ipl1–Sli15 at the centromere/inner kinetochore is dispensable for the establishment of chromosome biorientation

The engineered recruitment of Ipl1–Sli15 to the inner kinetochore with FRB-Sli15 and Mif2-FKBP12 (Figure 3A, iv) showed an equivalent rate of biorientation establishment to its physiological recruitment to the inner kinetochore/centromere (Figure 3A, i). Using this engineered recruitment, we next addressed whether a rapid turnover of Ipl1–Sli15 at the centromere/inner kinetochore is required to establish chromosome biorientation. In the presence of Rapamycin, FRB and FKBP12 bind each other so tightly that their interaction would be hardly exchanged or turned over [41]. In addition, the C-terminus of vertebrate INCENP shows extensive interaction around the ‘neck’ of Aurora B [42]. If this interaction mode is also conserved in yeast, Ipl1 and Sli15 may tightly interact with each other. Thus, Ipl1 and Sli15 may hardly be exchanged (or turned over) at the inner kinetochore in the engineered recruitment, but they can still support biorientation as efficiently as physiologically-recruited Ipl1–Sli15 (Figure 3A, i and iv).

Based on this surmise, we analyzed how rapidly Ipl1-GFP is turned over at the kinetochore in physiological recruitment and in engineered recruitment (using FRB-Sli15 and Mif2-FKBP12), using fluorescent recovery after photobleaching (FRAP). To distinguish localization of Ipl1-GFP at the kinetochore from that on spindle MTs, we isolated a chosen centromere, *CEN3*, from the mitotic spindle by inactivating *CEN3* and thereby inhibiting its kinetochore–MT interaction [37, 38], as shown in Figure 2A. Subsequently, we reactivated *CEN3* to allow kinetochore assembly and Ipl1-GFP localization on it (Figure 6A, i; 6B, i, 6C, i). We then photo-bleached Ipl1-GFP at the kinetochore (on *CEN3*) (Figure 6A, ii; 6B, ii; 6C ii) and analyzed recovery of the Ipl1-GFP signal on *CEN3* before *CEN3* reached the spindle following the interaction with a spindle MT (Figure 6A, iii). Immediately after photobleaching, the Ipl1-GFP signal was reduced to about 20% (relative to the signal before photobleaching; Figure 6B, ii; 6C, ii; 6D, 0 s). In physiological recruitment, Ipl1-GFP signals were rapidly recovered to 70– 80% within 30 s (Figure 6B, ii; 6D, blue line), suggesting a rapid turnover of Ipl1–Sli15 at the centromere/inner kinetochore. Subsequently (after 3 min), the Ip1-GFP signal on *CEN3* was gradually reduced, probably due to gradual photobleaching during image acquisition (Figure 6B, iii; 6D, blue line). By contrast, in engineered recruitment, the Ipl1-GFP signal hardly recovered on *CEN3* throughout observation up to 10 min (Figure 6C, ii, iii; 6D, red line), suggesting very little turnover of Ipl1–Sli15 at the inner kinetochore.

**Figure 6.**
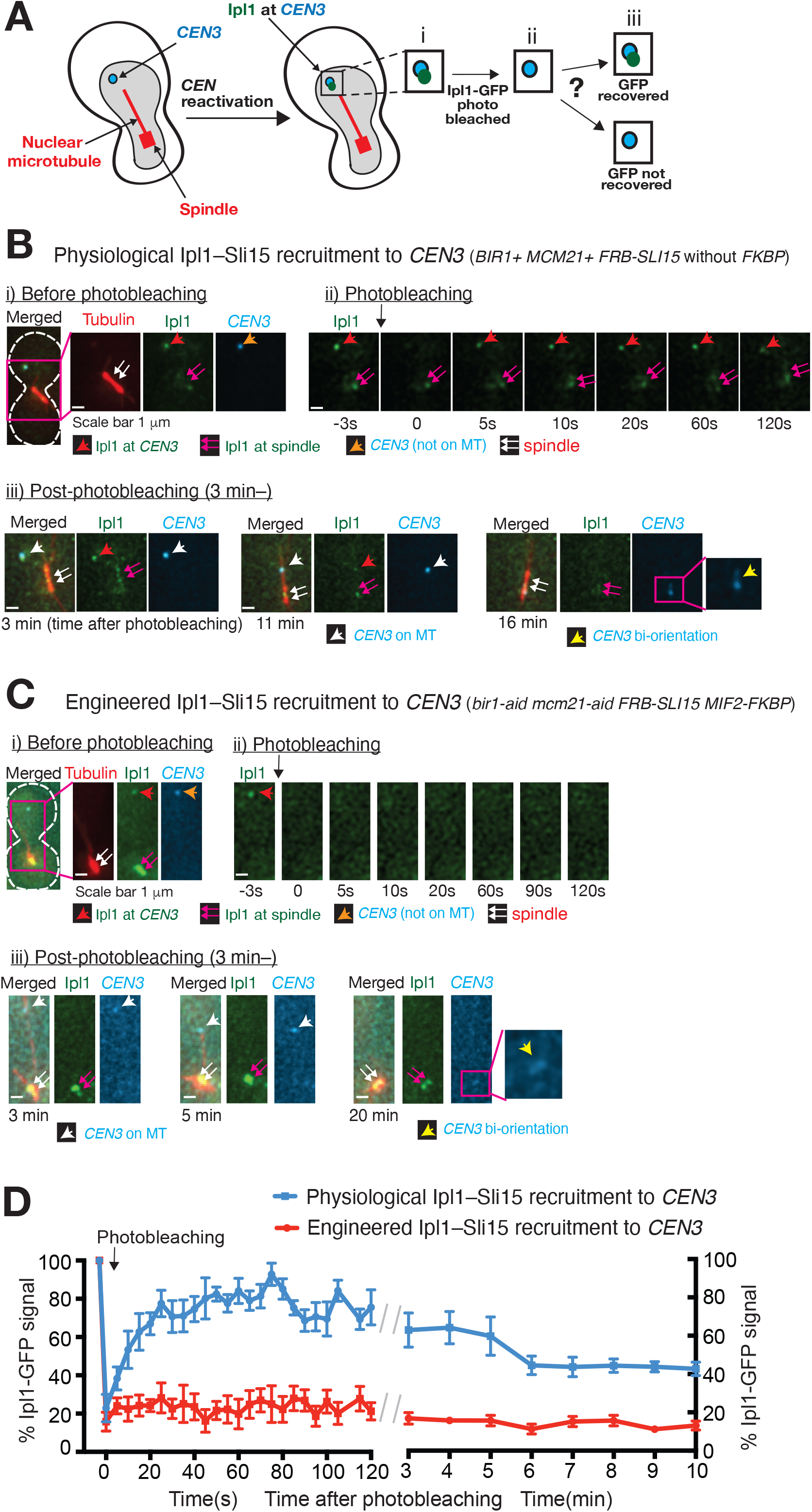
Ipl1–Sli15 shows rapid turnover at the centromere/inner kinetochore with physiological recruitment to it, but not with the engineered recruitment. **A**. Diagram shows the method of measuring Ipl1 turnover at *CEN3*/kinetochore using fluorescent recovery after photobleaching (FRAP) with the centromere reactivation system (see Figure 2A). Ipl1-GFP signal at the isolated and reactivated *CEN3* (and the kinetochore on it) was photobleached using 488 nm laser (i to ii). Subsequently, the recovery (or a lack of recovery) of the Ipl1-GFP signal was investigated (iii) to assess the turnover of Ipl1-GFP at *CEN3* (and the kinetochore on it). **B. C**. Representative outcomes of FRAP analyses in cells with physiological (**B**) and engineered (**C**) recruitment of Ipl1-Sli15 to *CEN3* (and the kinetochore on it). Two strains, *BIR1+ MCM21+* without *FKBP12* (T14015, **B**) and *bir1-aid mcm21-aid MIF2-2xFKBP* (T14024, **C**) carried common constructs *FRB-SLI15, IPL1-3xGFP, TIR, GAL1-10* promoter*-CEN3-tetOs, TetR-3xCFP, mCherry-TUB1* and *MET3* promoter*-CDC20*. Cells of the two strains were released from G1 arrest and subsequently arrested in metaphase by Cdc20 depletion while *CEN3* was inactivated. 500 µM NAA was added (to deplete Bir1-aid and Mcm21-aid if present) 30min before the release from G1 arrest, and 1 µM rapamycin was added (to induce FRB-FKBP12 dimerization if they are present) 30 mins before *CEN3* reactivation. After *CEN3* was reactivated during metaphase arrest, fluorescence images were acquired for GFP, CFP and mCherry on 9 z-sections (i). Then, GFP image was acquired on a single focal plane (ii, -3 s), followed by photobleaching of Ipl1-GFP signal at *CEN3* with 0.5 s pulse of 488 nm laser (from -0.5 to 0 s in ii). Immediately after this, a GFP image was acquired (0 s) on a single z section, followed by further image acquisition for 2 min with a 5-s interval (ii). Then GFP, CFP and mCherry images were acquired on 9 z-sections again, from 3 to 20 min (relative to photobleaching) with a 1-min interval (iii). For i) and iii), projected images (from 9 z-sections) are shown here. **D**. Following photobleaching, Ipl1-GFP signals rapidly recovered at *CEN3* with physiological recruitment, but not with engineered recruitment. In FRAP experiments described in **B** and **C**, Ipl1-GFP signals associated with *CEN3* were quantified along time in 6 cells of T14015 (physiological recruitment, blue line) and in 5 cells of T14024 (engineered recruitment, red line). Ipl1-GFP signals at *CEN3* were normalized to its pre-bleach signals (100%) in individual cells. Accordingly, the percentages of the Ipl1-GFP signal (mean and SEM at each time point) were plotted over time. Once *CEN3* reached the spindle (e.g. 16 min in **B**, iii; 20 min in **C**, iii), Ipl1-GFP signals could not be specifically measured at *CEN3* as other *CEN*s were also on the spindle, and therefore quantification stopped.

Nonetheless, the Ipl1–Sli15 at the inner kinetochore in engineered recruitment can support biorientation as well as Ipl1–Sli15 in physiological recruitment (Figures 3A, i and iv). We further tested this by investigating the kinetics of biorientation establishment after inactivation and subsequent reactivation of *CEN3* (Figure S2). Indeed, the engineered and physiological recruitments of Ipl1-Sli15 promoted biorientation establishment of sister *CEN3*s in very similar efficiency (Figure S2, i and ii). Altogether, these results imply that, although a rapid turnover of Ipl1–Sli15 occurs at the centromere/inner kinetochore in context of the physiological condition, this turnover is largely dispensable for biorientation establishment.

## Discussion

To test Aurora B spatial separation model and kinetochore conformational change model (see Introduction), we engineered recruitment of Ipl1–Sli15 to the inner kinetochore and outer kinetochore. In condition 1, the engineered recruitment was carried out in the absence of physiological Ipl1-Sli15 recruitment mechanisms (with depletion of Bir1 and Mcm21), prior to establishment of biorientation. In condition 2, the engineered recruitment was conducted in the presence of physiological Ipl1–Sli15 recruitment mechanisms (with intact Bir1 and Mcm21), after establishment of biorientation. Figure 1C shows the outcomes predicted from the above two models, in rectangles. We showed that the recruitment of Ipl1–Sli15 to the inner kinetochore a) restored biorientation establishment in condition 1 (Figure 3A), and b) did not disrupt biorientation maintenance in condition 2 (Figure 4A). These results are consistent with both models mentioned above (Figure 1C, green in rectangles). On the other hand, the recruitment of Ipl1–Sli15 to the outer kinetochore a) failed to restore biorientation establishment in condition 1 (Figure 3B), and b) disrupted biorientation maintenance in condition 2 (Figure 4B) relying on the Ipl1 kinase activity (Figure 5A). These results match predictions from the Aurora B spatial separation model (Figure 1C, red in rectangles), but not from the kinetochore conformational change model (Figure 1C, blue in rectangles). Collectively, our results suggest that 1) spatial separation of Aurora B from its outer kinetochore substrates is required to stabilize kinetochore–MT interaction when bi-orientation is established, and 2) kinetochore conformational change alone is insufficient to stabilize kinetochore–MT interaction.

However, while our results support the Aurora B spatial separation model, they do not exclude that other mechanisms are additionally required to stabilize kinetochore–MT interaction when bi-orientation is established. For example, a kinetochore conformational change may be additionally required. In fact, the evidence for the kinetochore conformational change model was recently obtained *in vitro* using purified yeast kinetochores [30]. If both Aurora B spatial separation mechanism and a kinetochore conformational change are required to stabilize kinetochore–MT interaction, the predicted outcomes in conditions 1 and 2 would be the same as the Aurora B spatial separation model alone (Figure 1C, red and green in rectangles), thus still matching our results in Figure 3–5.

Our results support the Aurora B spatial model in budding yeast. Given that Aurora B is crucial to promote error correction for biorientation from yeast to vertebrate cells, the Aurora B spatial separation model may also work in vertebrate cells. Previous studies showed that the ectopic targeting of CPC to the outer kinetochore destabilizes kinetochore–MT interaction [21, 43], which is consistent with the Aurora B spatial separation model. However, in contrast to budding yeast where CPC at both the centromere and inner kinetochore (Mcm21-Ctf19) is crucial for error correction [18, 19], it is still debated which fraction of the CPC within the centromere/kinetochore is important for error correction (i.e. destabilizes kinetochore–MT interaction with low tension) in vertebrate cells. For example, CPC at the centromere may be too far away from the kinetochore–MT interface to regulate this, due to mitotic chromosome condensation (which is negligible or modest in budding yeast) [44]. It was reported that a small fraction of CPC is present at the outer kinetochore in human cells and this fraction is diminished when tension is applied [45, 46] – if so, this reduction may be a key to stabilize kinetochore–MT interaction when biorientation is established. However, there are also reports that a fraction of CPC localizes at (or near to) the inner kinetochore [46, 47] – if this fraction is crucial for error correction, the Aurora B spatial separation mechanism may indeed work, i.e. stretched Ndc80C may separate Aurora B from its substrates (e.g. Ndc80 N-terminus) to stabilize kinetochore–MT interaction when bi-orientation is established and tension is applied. Further work is required to test the Aurora B spatial model in vertebrate cells.

Meanwhile, it is known that CPC shows rapid turnover on the centromere and kinetochore during early mitosis in mammalian cells [31-33]. We also observed such a rapid turnover of Ipl1–Sli15 on the centromere and kinetochore in budding yeast (Figure 6B, D; physiological recruitment). It was previously proposed that the turnover of CPC at the centromere/kinetochore is required for error correction to promote biorientation. For example, the turnover may facilitate Aurora B reaching its substrates in distance [20] and/or may promote shuttling of CPC between the centromere/kinetochore and the mitotic spindle to activate Aurora B kinase on spindle MTs [34]. We tested the importance of the rapid turnover of Ipl1–Sli15 in biorientation, using its engineered recruitment to the inner kinetochore. We showed that Ipl1–Sli15, recruited to the inner kinetochore by engineering, was hardly turned over (Figure 6C, D; engineered recruitment) and yet was able to support biorientation as efficiently as physiologically-recruited Ipl1–Sli15 (Figure 3A, S2). The corollary is that, in a physiological context, the rapid turnover of Ipl1–Sli15 happens at the centromere/inner kinetochore, but this is largely dispensable for error correction and biorientation, at least, in budding yeast. However, we still cannot rule out that the rapid turnover is required to enhance the fidelity of biorientation in yeast (even though not largely required for biorientation) or that the turnover of CPC at the centromere/kinetochore is essential for biorientation in vertebrate cells.

Collectively, our study gives important implications for how CPC promotes error correction for biorientation in a tension-dependent manner. Our results suggest that the spatial separation of Aurora B from its outer kinetochore substrates is required to stabilize kinetochore–MT interaction when tension is applied. Although Ipl1–Sli15 is rapidly turned over at the centromere/inner kinetochore, this turnover is not largely required for error correction and bi-orientation establishment. Error correction for chromosome biorientation lies at the heart of the chromosome segregation mechanism. Further study of biorientation in yeast and other organisms will elucidate mechanisms ensuring correct chromosome segregation in mitosis.

## Acknowledgement

We thank Tanaka lab members for discussion; S. Biggins, M. Kanemaki, U.K. Laemmli, K. Nasmyth, K. Bloom, R. Ciosk, K.E. Sawin, E. Schiebel, R.Y. Tsien, EUROSCARF and Yeast Resource Centre for providing reagents; and Dundee Imaging Facility for help in microscopy. This work was supported by the Wellcome Trust (219418/Z/19/Z).

## Author contribution

Conceptualization, T.U.T., L.J.G.-R. and S.L.; Methodology, L.J.G.-R. and S.L.; Investigation, S.L. and L.J.G.-R.; Formal Analysis, S.L. and L.J.G.-R.; Visualization; S.L., L.J.G.-R. and T.U.T., Writing; T.U.T., S.L. and L.J.G.-R., Supervision, T.U.T.; Project Administration, T.U.T.; Funding, T.U.T.

## Materials and Methods

### Yeast strain construction

All yeast *Saccharomyces cerevisiae* strains used in this study had a W303 background. The genotypes of the yeast strains are shown in Table S1. To make these yeast strains, precursor yeast strains were transformed with DNA constructs (see below) and/or two strains were crossed to each other followed by tetrad dissection and genotyping [48]. *IPL1* was tagged with *yEGFP* or *3xGFP* at its C-terminus at its original locus using *yEGFP-KanMX* cassette (pKT127 from EUROSCARF) or *3xGFP-KanMX* (pSM1023 from E. Schiebel’s lab) cassette as a PCR template, followed by yeast cell transformation with PCR products [49, 50]. *MIF2* and *NDC80* were tagged with 2x*FKBP* at their C-termini at their original loci using *2xFKBP-TRP1* and *2xFKBP-HI3* cassettes (P30582 and P30583 from EUROSCARF; Haruki, 2008 #143) as PCR templates followed by yeast cell transformation with PCR products. *BIR1* and *MCM21* were tagged with auxin-inducible degron (*aid*) at their C-termini at their original loci, using *IAA17-clonNAT* and *IAA17-spHIS5* cassettes (pT1450 and pT3656; *IAA17* [AID tag] was provided by Kanemaki’s lab [51]) as PCR templates, followed by yeast cell transformation with PCR products. To make FRB fusion to the N-terminus of Sli15, *SLI15* gene was replaced with *FRB-SLI15* at its original locus, using the *FRB-SLi15-ADE1-ADE2* construct flanked by *SLI15* non-coding regions (pT3655), for which *FRB* was provided by Laemmli’s lab [52] though EUROSCARF. The coding and non-coding regions of *ipl1-as5* allele (containing mutations M181G and T244A) were amplified by PCR from the *ipl1-as5* strain (SBY3056 from S. Biggin’s lab) and used to make *ipl1-as5-KanMX* construct flanked by *IPL1* non-coding regions (pT3676), which was subsequently used to replace *IPL1* wild type allele at its original locus. The constructions of *GAL1-10* promoter-*CEN3-tetOs, TetR-3xCFP, MET3* promoter-*CDC20, mCherry-TUB1, Spc42-4xmCherry*, and *CEN2-tetOs* were as previously described [18].

### Yeast cell culture

Methods for yeast cell culture were as previously described [48, 53]. To synchronize cells in the cell cycle, yeast cells were arrested in G1 phase by treatment with yeast mating hormone (α factor) and subsequently released to fresh media. Cells were cultured at 25°C in YPA medium (1% yeast extract, 2% peptone, 0.01% adenine hydrochloride) containing 2% glucose (YPAD), unless otherwise stated. To activate the *MET3* promoter, cells were incubated in a minus methionine drop-out medium. The *MET3* promoter was suppressed by adding 2 mM methionine to the relevant medium. To activate the *GAL1-10* promoter, cells were incubated in YPA medium containing 2% raffinose for at least 2.5 h, and subsequently, 2% galactose was added.

### Microscopy image acquisition and analyses

For live-cell imaging, yeast cells were immobilized either on a glass-bottom dish coated with concanavalin A or on a slide with an agarose pad and maintained either in synthetic-complete (SC) medium or in a mixture of SC plus YPA medium (3:1 ratio), as described previously [18, 37]. Images were acquired using DeltaVison Elite microscope (Applied Precision), a UPlanSApo 100× objective lens (Olympus; numerical aperture 1.40), SoftWoRX software (Applied Precision), and CoolSnap HQ2, Cascade II 512B EM-CCD or Pco-Edge 4.2 sCMOS camera (Photometrics) for image acquisition. Unless otherwise stated, we acquired nine z-sections (0.7 μm apart) at 25°C. Z-sections were subsequently deconvolved, projected to two-dimensional images and analyzed with Volocity software (Improvision). In Figure 3–5, it was judged that biorientation was established or maintained if sister *CEN2* fluorescent dots showed either a) complete separation or b) stretching in the middle of the spindle, for more than 6 imaging time points (out of 10 time points).

### FRB-FKBP12 dimerization and depletion of an AID-tagged protein

FRB-FKBP12 dimerization was induced by the addition of 1 μM rapamycin to the culture medium. To keep yeast cells alive in the presence of rapamycin and to make FRB-FKBP12 dimerization efficient, yeast strains also carried *TOR1-1* and *Δfpr1* [52]. Meanwhile, to induce depletion of AID-tagged proteins, 0.5 mM auxin NAA (1-naphaleneacetic acid) was added to the culture medium. To facilitate the depletion of an AID-tagged protein, yeast strains also carried *TIR1* gene of rice *Oryza sativa* (*osTIR1*), whose expression was driven by *ADH1* promoter [51].

### Centromere reactivation assay

To analyze localization and dynamics (with FRAP) of Ipl1 at the centromere isolated from the spindle, the centromere re-activation assay was used [37, 38]. In this assay, kinetochore assembly was delayed on a chosen centromere by transcription from the *GAL* promoter (*GAL1-10* promoter-*CEN3-tetOs* replacing *CEN15* on chromosome XV). This increased the distance between the centromere and the mitotic spindle during metaphase arrest, allowing observation of protein localization specifically at *CEN3* after inducing kinetochore assembly on the centromere by turning off the *GAL* promoter in metaphase-arrested cells. Cells with *GAL1-10* promoter-*CEN3-tetOs* and *MET3* promoter-*CDC20* were cultured overnight in methionine drop-out media with 2% raffinose, treated with α factor for 2.5 hours (to arrest cells in G1 phase), and released to fresh media with 2% raffinose, 2% galactose and 2 mM methionine (for Cdc20 depletion and *CEN3* inactivation). After 2 hours, cells were suspended in an SC medium containing 2% glucose and methionine to reactivate *CEN3*. Protein localization was analyzed at *CEN3* after *CEN3* reactivation and before *CEN3* interaction with microtubules extending from the spindle. After z-sections were projected to 2D images, Ipl1-GFP and Sli15-FRB-GFP signals (associated with *CEN3*) were quantified in the 2×2 pixel area at their maximum intensity, using the Voxel Spy tools of Volocity (Improvision). The intracellular background was subtracted in each measurement.

### Fluorescence recovery after photo-bleaching (FRAP) assay

The FRAP experiments were carried out on a DeltaVision Elite microscope with the Quantitative Laser Module (Applied Precision) and Pco-Edge 4.2 sCMOS camera (Photometrics). FRAP experiments were performed at 25ºC in three successive steps as follows: 1) pre-bleach: a pre-bleach fluorescence image was acquired in GFP, CFP and mCherry channels with 9 z-sections (0.7 μm apart); 2) photo-bleach: a single z-section image of GFP was acquired with a 488 nm laser line at a low laser intensity (at -3 s). Photobleaching was carried out by a 0.5-s pulse of the 488 nm laser (from -0.5 s to 0 s) at 75% laser power using a point bleach tool to target the Ipl1-GFP signal at *CEN3* that is distanced from the spindle (see centromere reactivation assay). Immediate after photobleaching (at 0 s) and thereafter with 5-s intervals for 2 min, a single z-section image of GFP was acquired; 3) post-bleach: post-bleach fluorescence images were acquired in GFP, CFP and mCherry channels with 9 z-sections with 1-min intervals for 20 min. The Ipl1-GFP signal at *CEN3* was quantified until *CEN3* reached the spindle, as described above. Note that, in Figure 6C ii), we interpret that Ipl1-GFP signals did not recover after photo-bleaching, rather than they became out of focus, for the following reasons – a) it is very unlikely that, during 5–120 s, Ipl1-GFP signals recovered at *CEN3* but became out of focus in the single z-section images, because Ipl1-GFP signals were not detected even if *CEN3* was detected in the 9 z-section images at 3 and 11 min in Figure 6C iii), and b) in our analysis of physiologically-recruited Ipl1-GFP that was photobleached and subsequently recovered (Figure 6B ii), it was rare Ipl1-GFP signals at *CEN3* became out of focus during 5–120 s in single z-section images with 5-s intervals.

### Statistical analysis

Statistical analyses were carried out using Prism software (GraphPad). Methods of statistic tests are stated in figure legends. t-test was used in Figures 2 and S1 to determine whether GFP signals are significantly different between groups. Fisher’s extract test was used in Figures 3, 4 and 5 to address if there is a significant difference in the frequency of biorientation. The null hypotheses in these tests were that the samples were collected randomly and independently from the same population. All *p* values were two-tailed, and the null hypotheses were reasonably discarded when *p* values were smaller than 0.05.

## Figures and Supplementary materials

### Supplementary Figures

**Figure S1.**
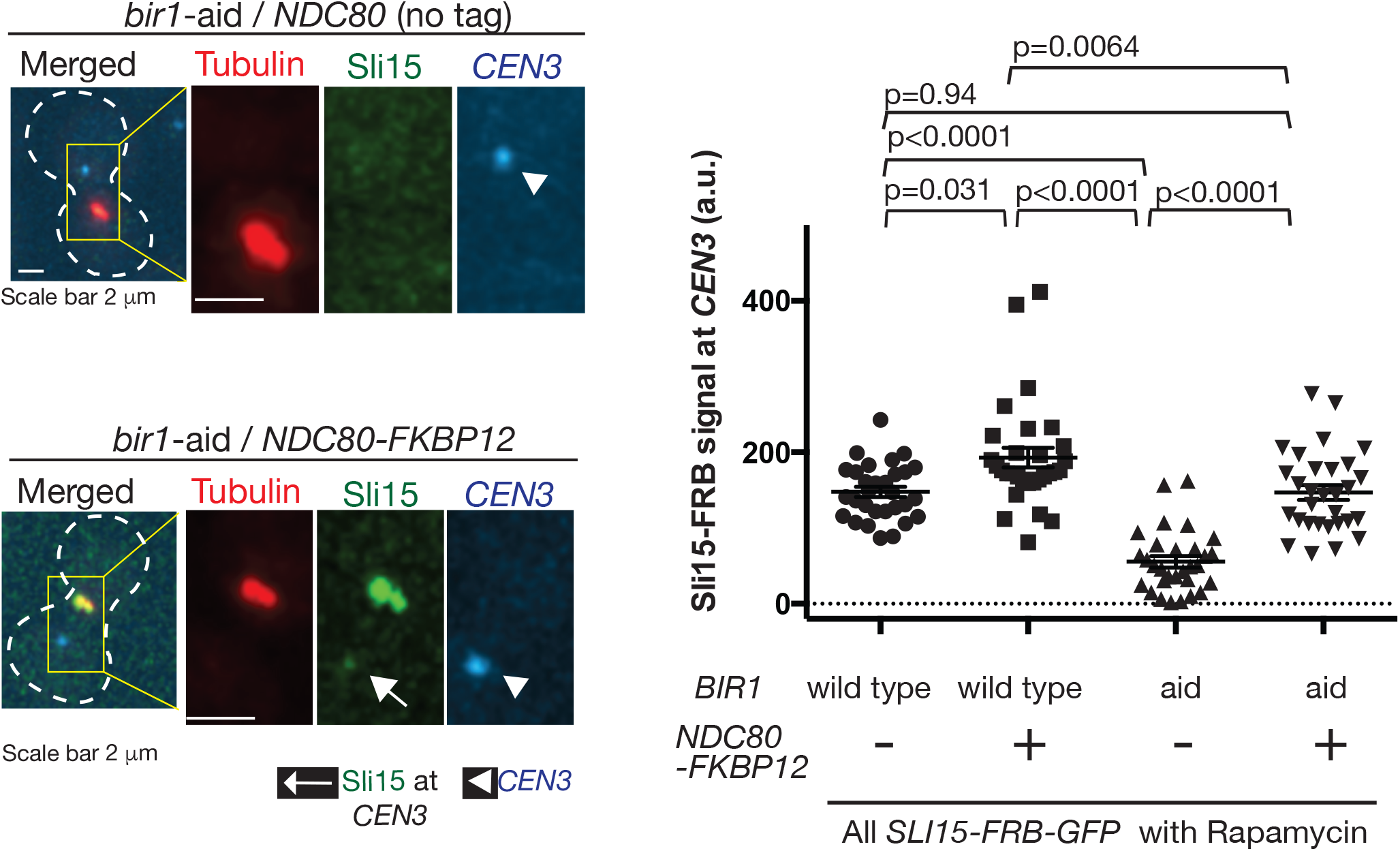
The engineered Ipl1–Sli15 recruitment to the outer kinetochore in the absence of Bir1 restores its physiological level at the kinetochore using FRB fusion to the C-terminus of Sli15. Four strains contained the combination of i) *BIR1+* (wile-type) or *bir1-aid* and ii) with or without *FKBP* fusion to *NDC80* (*NDC80-FKBP12*), as shown below in graph (T13999, T13800, T13201 and T13803 from left to right). All four strains also carried *SLI15-FRB-GFP* (fusion of FRB-GFP to the C-terminus of Sli15), *TIR, GAL1-10* promoter*-CEN3-tetOs, TetR-3xCFP, mCherry-TUB1* and *MET3* promoter*-CDC20*. Cells were released from G1 arrest and subsequently arrested in metaphase by Cdc20 depletion while *CEN3* was inactivated. Immediately after *CEN3* was reactivated during metaphase arrest, fluorescence images were acquired for 10 min with a 1-min interval. 500 µM NAA was added (to deplete Bir1-aid if present) 30min before the release from G1 arrest, and 1 µM rapamycin was added (to induce FRB-FKBP12 dimerization if they are present) 30 mins before *CEN3* reactivation. Representative cells (T13201 [top] and T13803 [bottom]) are shown on the left. Cell shapes are shown in white broken lines. The Sli15-FRB-GFP signals associated with *CEN3* were quantified as in Figure 2D, and shown in graph (right), with mean and error bars (SEM). *p* values were obtained by t-test.

**Figure S2.**
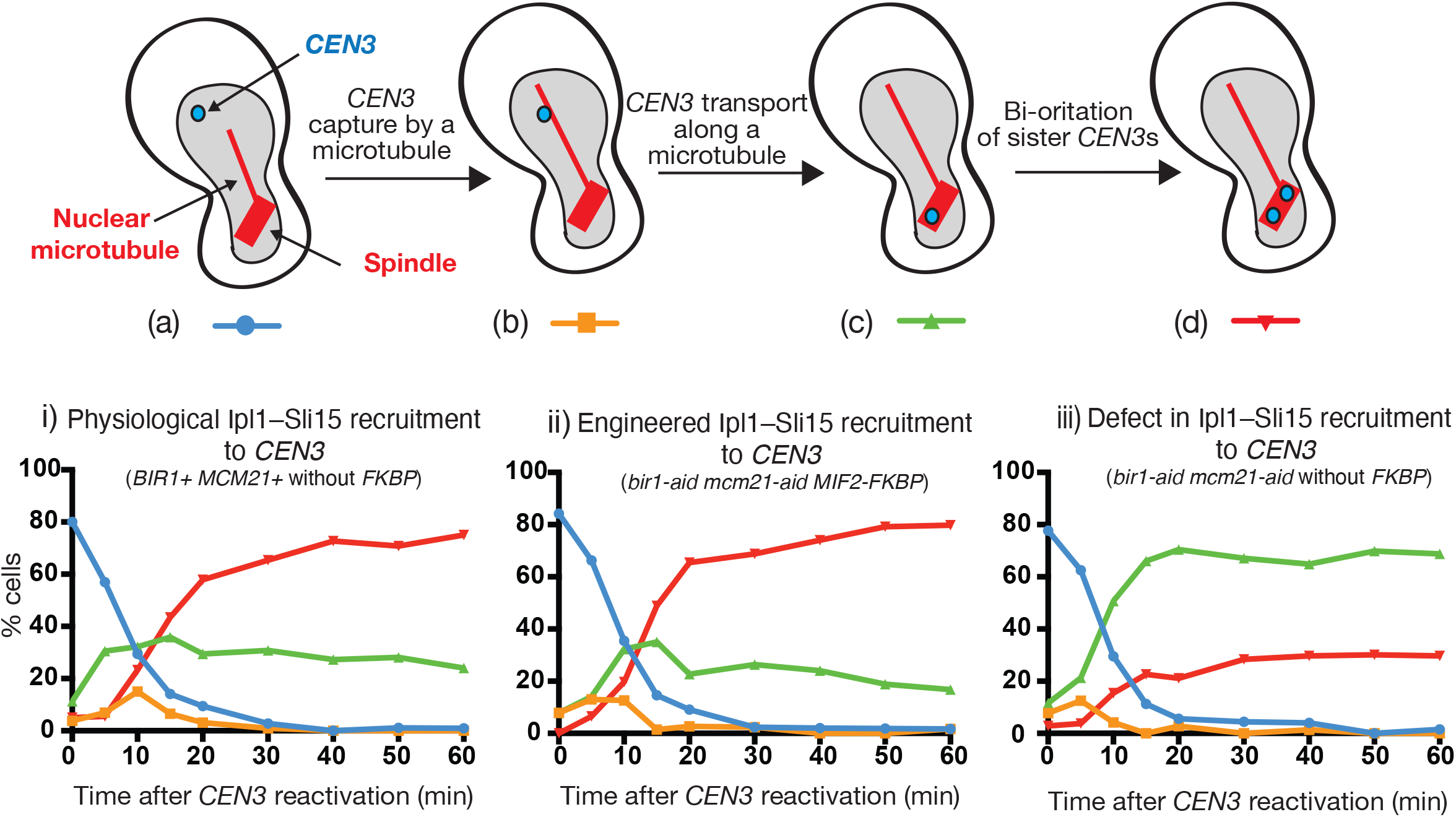
Physiological and engineered recruitments of Ipl1–Sli15 to the centromere/inner kinetochore promote the establishment of chromosome biorientation with similar efficiency after *CEN3* reactivation. Three strains, *BIR1+ MCM21+* without *FKBP12* (T14015, i) and *bir1-aid mcm21-aid MIF2-2xFKBP12* (T14024, ii), and *bir1-aid mcm21-aid* without *FKBP12* (T14022, iii) carried common constructs *FRB-SLI15, IPL1-3xGFP, TIR, GAL1-10* promoter*-CEN3-tetOs, TetR-3xCFP, mCherry-TUB1* and *MET3* promoter*-CDC20*. These strains were used to represent physiological Ipl1–Sli15 recruitment to the centromere/inner kinetochore (i), engineered Ipl1– Sli15 recruitment to the inner kinetochore (ii), and defect in Ipl1–Sli15 recruitment to the centromere/inner kinetochore (iii). Cells of the three strains were released from G1 arrest and subsequently arrested in metaphase by Cdc20 depletion while *CEN3* was inactivated. 500 µM NAA was added (to deplete Bir1-aid and Mcm21-aid if present) 30min before the release from G1 arrest, and 1 µM rapamycin was added (to induce FRB-FKBP12 dimerization if they are present) 30 mins before *CEN3* reactivation. After *CEN3* was reactivated during metaphase arrest, aliquots of cells were collected and fluorescence images were acquired at 0, 5, 10, 15, 20, 30, 40, 50, and 60 min. At each time point, the position of *CEN3* was categorized, as shown in diagrams (top); (a) *CEN3* has not yet been captured by a MT (blue line), (b) *CEN3* is on a MT extending for a spindle pole (orange line), (c) *CEN3* is on the spindle, but sister *CEN3*s are not separated (green line). (d) Sister *CEN3*s are separated on the spindle, i.e. biorientation has been established (red line). The percentage of cells with *CEN3* in each category was plotted in the graphs (bottom; 53–146 cells were evaluated in total at each time point). The results show that 1) the timing of *CEN3* capture by a MT is similar between the three conditions (compare blue lines in i, ii and iii), 2) engineered recruitment of Ipl1–Sli15 to the inner kinetochore promotes the efficient establishment of biorientation (compare green and red lines in ii and iii), and 3) the efficiency of biorientation is similar between physiological and engineered recruitments of Ipl1–Sli15 to the centromere/inner kinetochore (compare red lines in i and ii).

**Supplemental Table S1.**
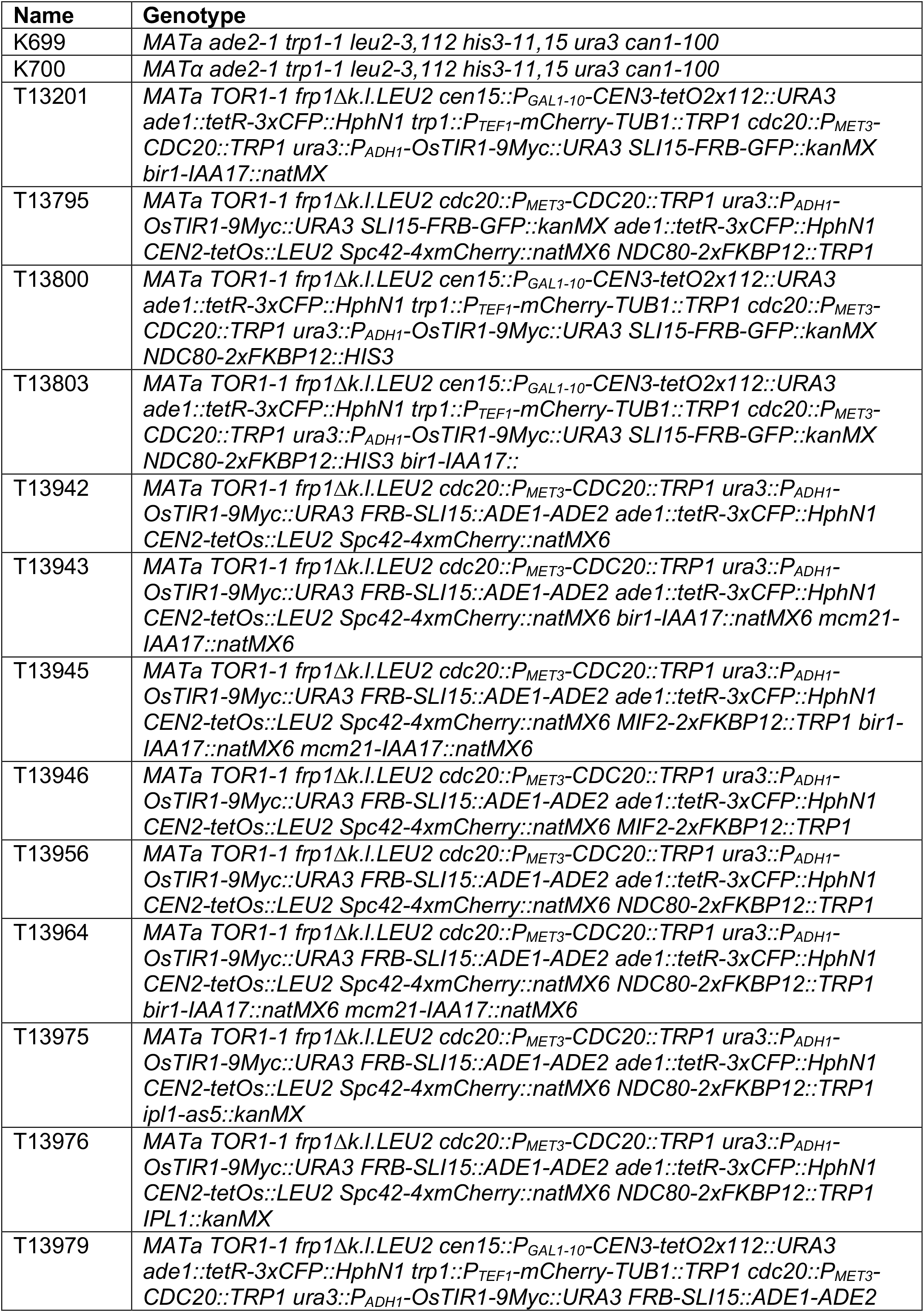
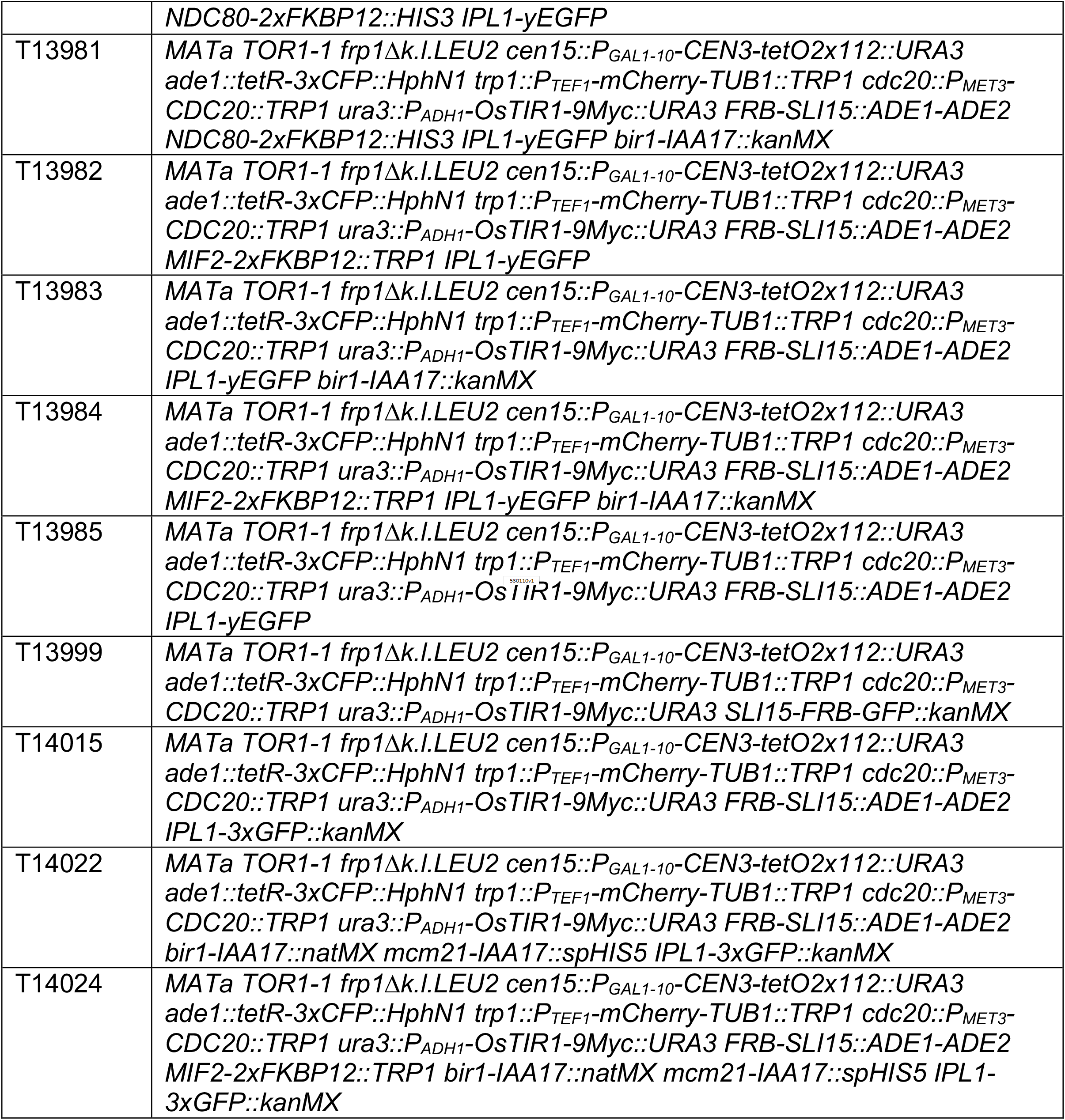
The table shows genotypes of yeast strains used in this study. All strains used in this study are derivatives of *Saccharomyces cerevisiae* W303 (K699 and K700 from K. Nasmyth lab).

